# Specificity of Aberrant Exons in a Saturated Background of Differentially Expressed Exons

**DOI:** 10.1101/2025.06.25.661610

**Authors:** Urmi Das, Hai Nguyen, Jian Yang, Samuel Ogunsola, Madeleine Hamilton, Yujia Wu, Ling Liu, Wenguang Cao, Chunyu Liu, Jiuyong Xie

## Abstract

Determining the maximal specificity of aberrant exons from multifactorial disease tissues in a whole body background could provide insights into the exon origin as well as potential targets of diagnosis or therapy, but it remains a challenging task. For this purpose, we obtained a saturated list of differentially expressed exons (DEXs) from our independent reference human exome extracted from 56 normal tissues by DEXSeq. We found that the DEXs comprised the majority of the reference exome and that 99.4% of cancer-specific exons relative to paired normal tissues fell within the DEX list, consistent with the ectopic expression of the majority of aberrant exons. We then screened over seven thousand pathogenic single nucleotide variants (SNVs) in the TCGA and COSMIC databases by SpliceAI. The analysis identified over three hundreds of highly confident, mostly somatic SNV-specific splicing events and/or novel exonic fragments in five types of adenocarcinomas. Interestingly, these events are all associated with *cis*-acting mutations in tumor suppressor genes, particularly in *TP53* with an antigenic novel peptide. This revelation of the DEX exome dominance and non-specificity of nearly all aberrant exons not only reshapes our understanding of the normal human exome by highlighting its mainly variable over constitutive exons, but also illuminates the path through *cis*-acting somatic and pathogenic SNVs to identify genuine cancer-specific exonic fragments that are absent in any normal tissues, beyond neo-exon junctions. This path holds promise for identifying novel peptide fragments for antigens in holistic cancer treatments including vaccination and immunotherapies.

## Introduction

For the maximal specificity of aberrant exons, their expression would ideally be limited to the diseased tissue of a patient, avoiding any potential undesired effects in clinical applications. However, about 80% of aberrant exons in cancer were found in over 30 normal tissues, at least according to exon-exon junction studies of alternatively spliced exons^1,2^. Besides alternative splicing (AS), alternative transcription start (ATS) or polyadenylation (APA) could also produce aberrant exons^1,3-6^, but the specificity of aberrant exons overall in a whole body background remains unknown.

Differentially expressed exons (DEXs) from the AS, ATS or APA contribute greatly to the transcriptomic and proteomic diversity in metazoans^7-11^. While alternative splicing has been found in over 90% of human genes^12,13^, the proportion of alternatively spliced exons and DEXs in general in the human exome — which comprises all unique exons in the genome — is not yet determined. As such, the percentage of constitutive exons is undetermined as well. Determining the prevalence and saturated number of DEXs will not only present a more refined view of the diverse human transcriptome but also establish a broader context for assessing the specificity of exons in a background closer to the whole body exome, as potential targets of cancer or other disease therapy.

Research has consistently shown dysregulation in the transcriptome across various cancer types with aberrant AS, ATS, and APA ^1,3-6^. For example, tumors display 30% more alternative splicing events than the corresponding normal tissues ^1^. Key players behind this include mutations in *cis*-acting elements or *trans*-acting factors. These mutations can disrupt or modify canonical splice sites leading to aberrant splicing, and can even create new splice sites ^14,15^. Notably, in deep introns, around 16% of tumor samples from various types show a multitude of splice site-creating mutations ^16^. Moreover, as cancer progresses, mutations accumulate. The number and frequency of these mutations can vary widely across genes, types of cancer, and even among patients ^17^. The diverse nature of these mutations and their impacts on RNA processing suggest a vast array of aberrant exons present in tumor cells. While the specificity of these exons to cancer has primarily been studied in comparison to their respective tissues of tumor origin (e.g., breast cancer versus normal breast tissue) or neo-exon junctions over 30 normal tissues^1,2^, aberrant exon specificity relative to a saturated DEX exome remains unclear.

In this study, we analyzed the exome of 56 human tissues to identify a saturated list of DEXs and assessed exon specificity in cancers. Our findings reveal that nearly all cancer-specific exons by paired normal tissue comparisons are DEXs within a DEX-dominated normal human exome. We show that the truly cancer-specific ones in this exome background are those associated with somatic single nucleotide variants (SNVs) with potential to promote the generation of novel peptides and cancer-specific antigens.

## Results

### 1. DEXs dominate the human exome as the majority of exons

To estimate the total number of differentially expressed exons (DEXs) in the human exome, we examined the RNA-Seq data of 56 nominally normal human tissues (53 adult plus 3 fetal tissues) ^18-20^. Data were obtained from the NCBI SRA and dbGaP databases, excluding cell lines and disease samples where aberrantly processed variants are often found ^21,22^. From a total of 12.7 billion of paired-end raw reads, we successfully mapped 8.8 billion of high-quality concordant pairs to the transcriptome of Ensembl GRCh38.78 (about ∼157 millions of reads per tissue on average, S_Fig. 1a). To analyze differential exon usage between a tissue pair by DEXSeq ^23^, the Ensembl-annotated exons in the genome/transcriptome were divided into unique bins (hereafter called exons for simplicity) based on the exon boundaries among different transcripts. In total, 370,559 exons from 27,463 genes (17,655 protein-coding) with a minimum average of 20 reads per base (exon base mean, EBM) are included in the reference exome for subsequent analyses (Fig. 1A).

**Figure 1.**
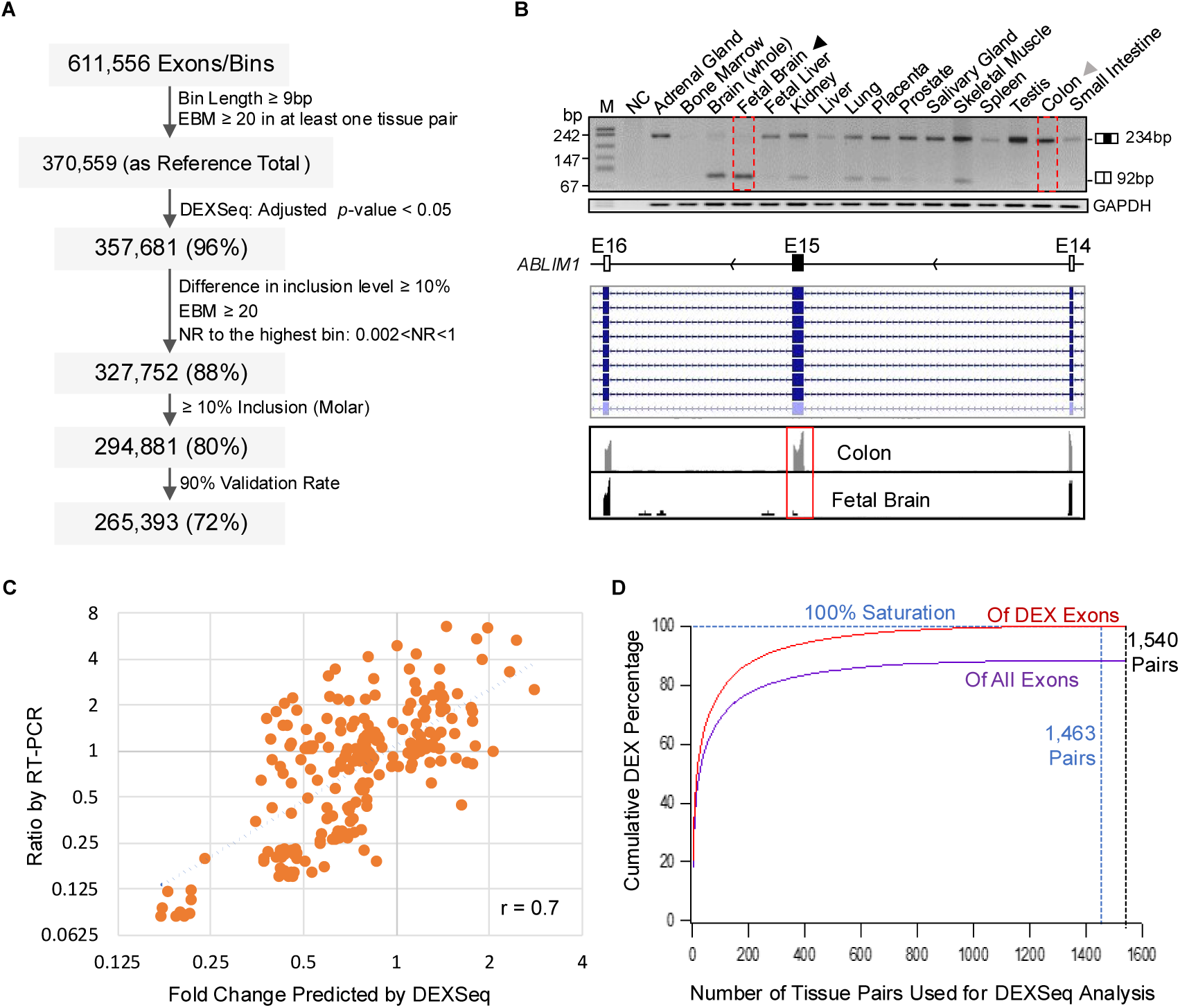
Estimating the saturating proportion of DEXs in the human exome. **A.** Flowchart of our filtering pipeline for differential exon usage analyses in 56 types of nominally normal human tissues. NR: ratio of normalized reads. **B.** Validation of DEXs by semi-quantitative RT-PCR. An example illustrating our validation of predicted alternative exons: predicted differential usage of the *ABLIM1* exon 15 (ENST00000533213.6), a curated constitutive exon among the RefSeq alternate haplotypes in the UCSC Genome (Upper panel), between the fetal brain and adult colon tissues in the IGV (Middle panel) and an agarose gel of RT-PCR products from 16 different types of human tissues (Lower panel). M: marker, NC: PCR negative control, GAPDH: RNA loading control. **C**. Correlation of predicted fold change (X-axis) with the ratio of exon inclusion percentages of RT-PCR products (Y-axis) between samples in log2 scales (n = 240 alternative splicing events of 12 genes in 110 tissue pairs). **D.** Saturation curve of the DEXs by their percentages of DEXs only (n = 327,752) or the total number of exons (n = 370,559) in the exome of 56 types of human normal tissues.

Interestingly, 357,681 exons, comprising 96% of the reference exome, exhibited significant differences (adjusted *p* < 0.05) in normalized read counts in at least one of the 1,540 tissue pairs. More stringent filtering with a RT-PCR-validated protocol as we have reported ^24^, resulted in 88% DEXs (327,752 exons, Fig. 1A). Increasing the EBM from 20 up to 1000 obtained similar proportions (S_Fig. 1b, 88.57% ± 0.83%, mean ± SEM, n = 14 thresholds of EBMs). At lower cut-off *p*-values (down to <0.0001), higher EBM numbers accompanied higher DEX abundance. Together, these data support that the 88% of DEX is accurate but not an overestimate at the *p* value < 0.05 or EBM ≥ 20.

After further examination of a group of exons in the Integrative Genomics Viewer (IGV) ^25^, we designed primers and successfully amplified over 200 alternative splicing events in semi-quantitative RT-PCR (Fig. 1B & C). The resulting exon inclusion ratio between tissue samples correlates with the DEXSeq prediction at a Pearson coefficient 0.7 (Fig. 1C), with a DEX validation rate of 90% (Fig. 1A). The RT-PCR data also allowed us to estimate that about 90% of the DEXs (294,881 exons) are included in at least 10% (in molar ratio) of their gene transcripts, comprising at least 79% of the reference exome. Furthermore, of the 349,833 annotated non-overlapping human exons in Ensembl release 93, only 15,990 (4.5%) have the same paired chromosome start-end coordinates in all annotated transcripts, likely as constitutive exons. Therefore, it is likely that the majority (> 79%) of human exons are DEXs rather than constitutive exons.

To determine whether the DEX percentage of 79% has reached the saturation level, we plotted the accumulating number of unique DEXs against the corresponding tissue pairs (Fig. 1D). The number of DEX increased exponentially with the addition of tissue pairs, reaching 90% of the reference total at 252 pairs, 99% at 868 pairs, and 100% at 1463 pairs without further increase from the additional 77 pairs. Thus, the pairwise search has saturated the DEXs in the 56-tissue exome.

To determine the relative contribution by AS-, ATS-or APA-mediated differential exon usage, DEXs were tagged as terminal or internal exons following transcriptome annotation. We identified 127,697 internal, 87,521 terminal and 79,663 as either internal or terminal DEXs depending on the transcript (S_Fig. 2a). By random examination of three subsets of DEXs (n = 100 each), we estimated that 43% were mediated by AS and 57% by ATS or APA (S_Fig. 2b). In particular, the ATS/APA-mediated DEXs fell into two groups, either in a longer terminal exon with DEXs from ATS or APA sites or internal exons between alternative terminal exons of different transcripts of the same host gene. The latter comprises 53% of all ATS/APA-induced DEXs.

**Figure 2.**
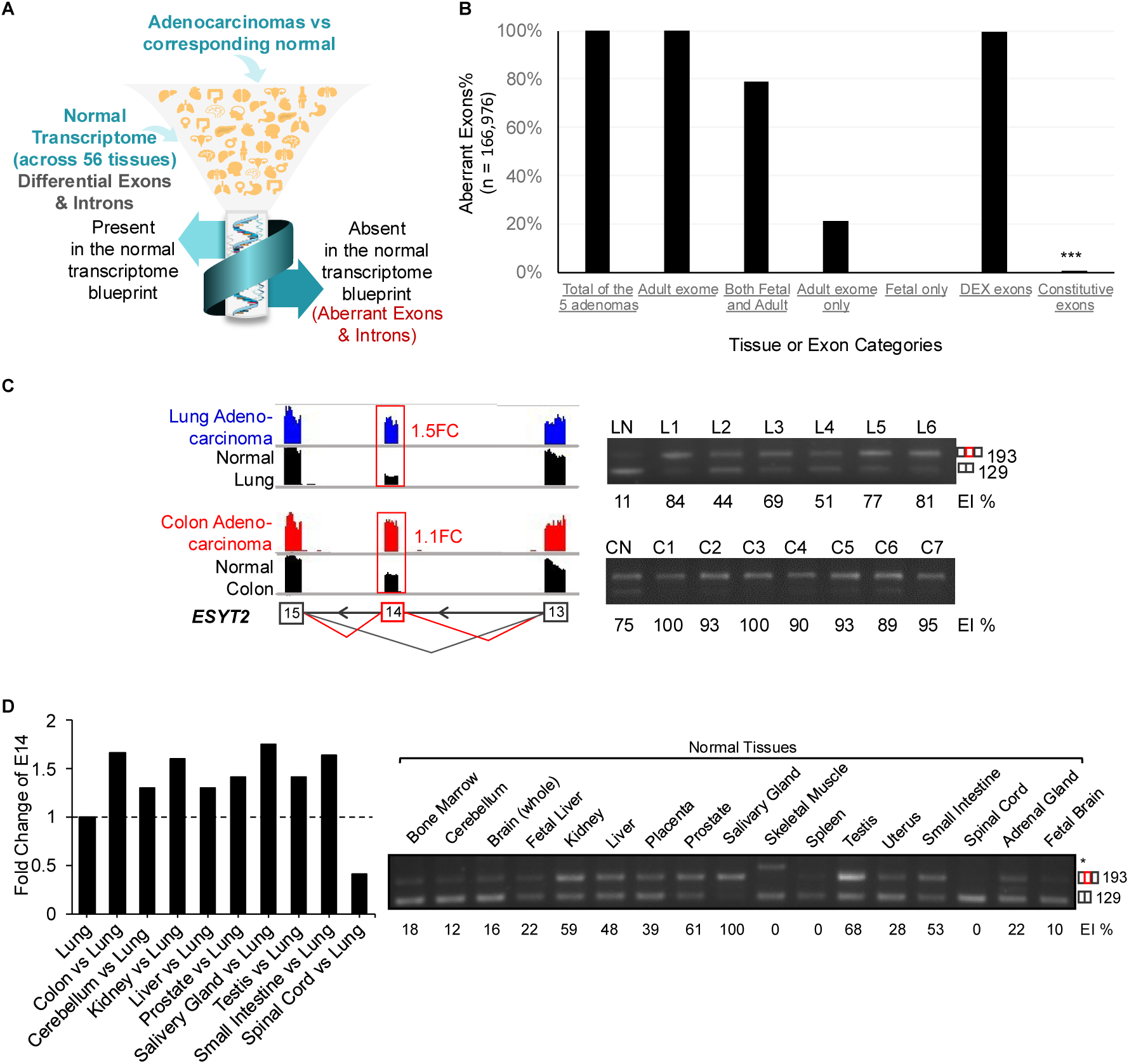
Ectopic expression of aberrant exons in adenocarcinoma. **A**. An overview of cross-checking the aberrations in adenocarcinoma against the normal transcriptome. **B**. Bar graph of the percent distribution of the aberrant exons among the adult or fetal, and among the DEX or constitutive exons. **C**. IGV view of E14 of *ESYT2* gene in lung and colon adenocarcinoma with fold change predicted by DEXSeq and validation by semiquantitative RT-PCR in lung and colon adenocarcinoma tumor samples (Upper panel). **D**. Fold change of E14 in other normal tissues compared to normal lung by DEXSeq analysis; and validation of the differential exon usage of E14 in 15 different ‘normal’ adult tissues and 2 fetal tissues (Lower panel). EI%-Exon inclusion percentage, M-Marker, NC-Negative control, LN-Normal Lung, (L1-L6) 6 Lung adenocarcinoma patients, CN-Normal colon, (C1-C7) 7 Colon adenocarcinoma patients.

To determine the DEX distribution profile across tissues, we performed pair-wise symmetric correlation based on the number of DEXs per tissue-pair. The brain regions, when compared with other tissues including duodenum, urinary bladder, appendix, kidney, esophagus, and endometrium, showed much higher variabilities yielding more DEXs, than among the brain regions themselves (S_Fig. 3). Higher variabilities were also observed between the fetal and adult tissues.

**Figure 3.**
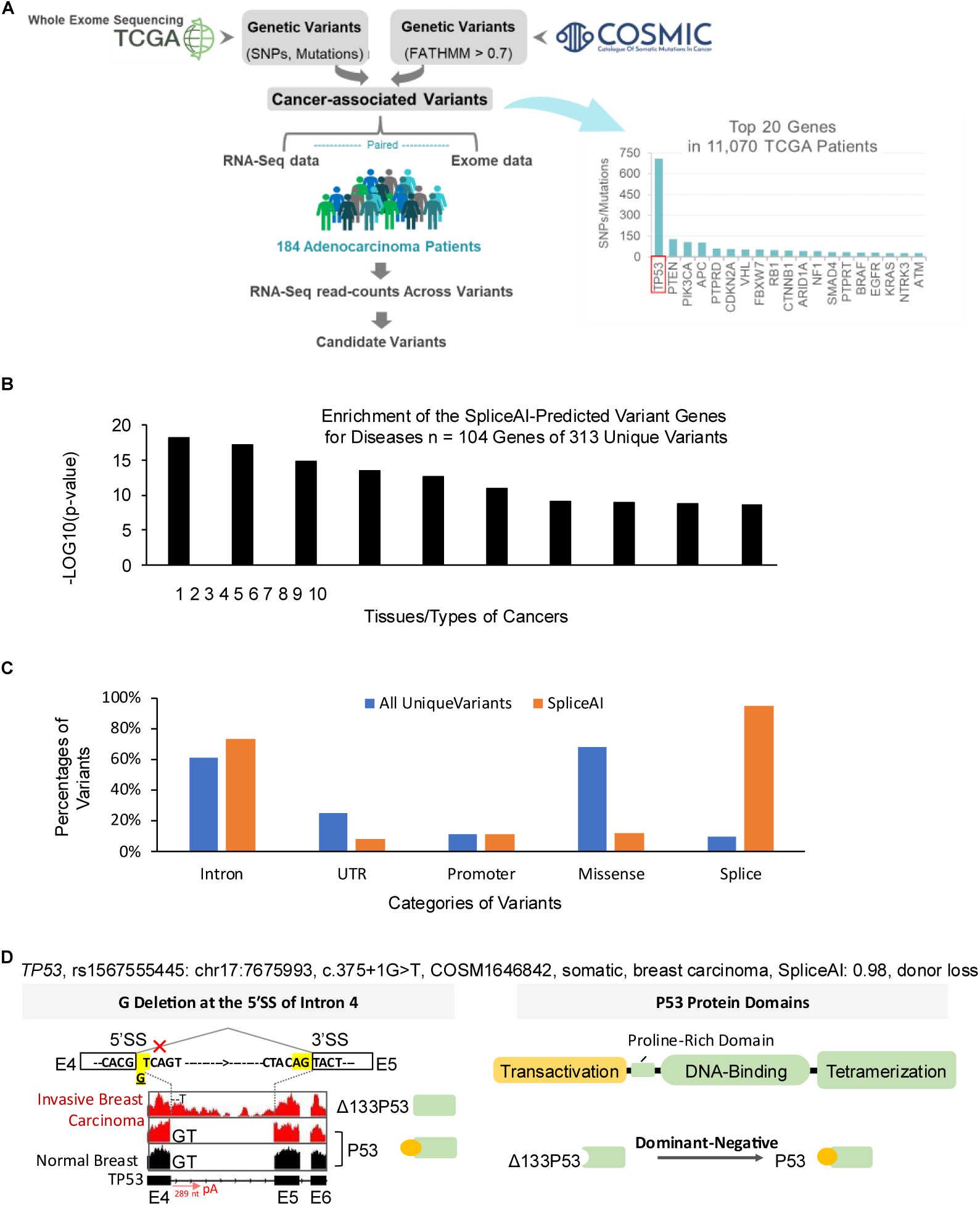
Identifying pathogenic SNVs mediating authentic cancer-specific aberrant exons by SpliceAI. **A.** An overview of filtering process pathogenic genetic variants from WGS to verify mediated splicing aberrations by the paired RNA-seq data. **B.** Top 10 DISGENET disease clusters of the 104 genes of 313 unique variants with SpliceAI scores > 0.8, identified by SpliceAI from a starting total of 7,085 unique variants with pathogenic FATHMM scores >0.5 in both the TCGA and COSMIC databases. The diseases are: 1. Breast adenocarcinoma, 2. Leukemia, Myelocytic, Acute, 3. Ovarian Carcinoma, 4. Conventional (Clear Cell) Renal Cell Carcinoma, 5. Bladder Neoplasm, 6. Malignant neoplasm of pancreas, 7. Adenocarcinoma, 8. Squamous cell carcinoma of esophagus, 9. Cholangiocarcinoma, 10. Malignant neoplasm of stomach. **C**. Distribution of unique variants in the starting pool and the SpliceAI > 0.8 group. **D**. Novel exon fragments generated after the abolishment of canonical splice site by a SNV. The IGV view of *TP53* in breast carcinoma patient with (top panel) or without (middle panel) delG in the 5′SS. Deletion of G in 5′SS of exon 4 likely lead to the use of ATS at exon 5 (NM_001407266.1) or intron retention. The resulting transcripts, when translated, generate N-terminally truncated p53 proteins lacking the transactivation domain. On the bottom, is a diagram of the domains of protein p53. P53 without transactivation domain acts as dominant negative inhibitor of its wild type.

### 2. Nearly all aberrant exons of adenocarcinomas belong to the saturated DEX exome

We then selected adenocarcinomas of five organs, lung, colon, pancreas, prostate, and breast by their worldwide high incidence and mortality^26^, and analyzed their RNA-Seq data of 184 patients from the Cancer Genome Atlas (TCGA) database ^27,28^. The samples were compared with their paired normal tissues using DEXSeq, to identify 167,015 unique, differentially used exons of 13,638 genes with at least 1.1-fold change over normal samples (adjusted *p* < 0.005). We found that 99.8% of them were among the 370,559 exons of the normal reference exome (Fig. 2A), of which 99.9% in the adult and 78.9% in the three fetal tissues (Fig. 2B), suggesting that nearly all aberrant exons in cancers are also normally expressed in the other tissues. Therefore, it is likely that the majority of aberrant exons are expressed ectopically in cancers.

Of these aberrant exons, 21% were found to be adult tissue-specific and 79% were found in both the adult and fetal tissues (Fig. 2B), representing a 14-fold enrichment in the fetal over the adult stage per tissue (*p* = 1.5E-30, hypergeometric density test). This observation supports the activation of fetal stage splicing or RNA processing programs in cancers. However, there was no fetal-specific exons, suggesting that all these aberrant exons in cancers are also expressed in at least one of the adult tissues as alternative exons.

Moreover, 99.4% of the aberrant exons were among the list of 327,752 DEXs and about 0.5% among the constitutive exons. They comprised of 51% and 2% of the DEX and constitutive exons, respectively, a 27-fold enrichment among the DEX exons (*p* = 0, hypergeometric density test). We validated a group of the aberrant exons and their ectopic expression by RT-PCR of tumor and normal samples (S_Figs. 4-5). For instance, the exon 14 (E14) usage in the *EYST2* (Extended synaptotagmin-2) transcripts increased not only in the lung and colon adenocarcinomas but also in the normal salivary gland, colon, kidney and prostate over the normal lung (Fig. 2C-D). Similar ectopic expression was found for the E3b of *RAC1* (Rac Family Small GTPase 1, S_Fig. 5). Therefore, it is likely that these aberrant exons in cancers are nearly all in adult tissues as alternative exons.

**Figure 4.**
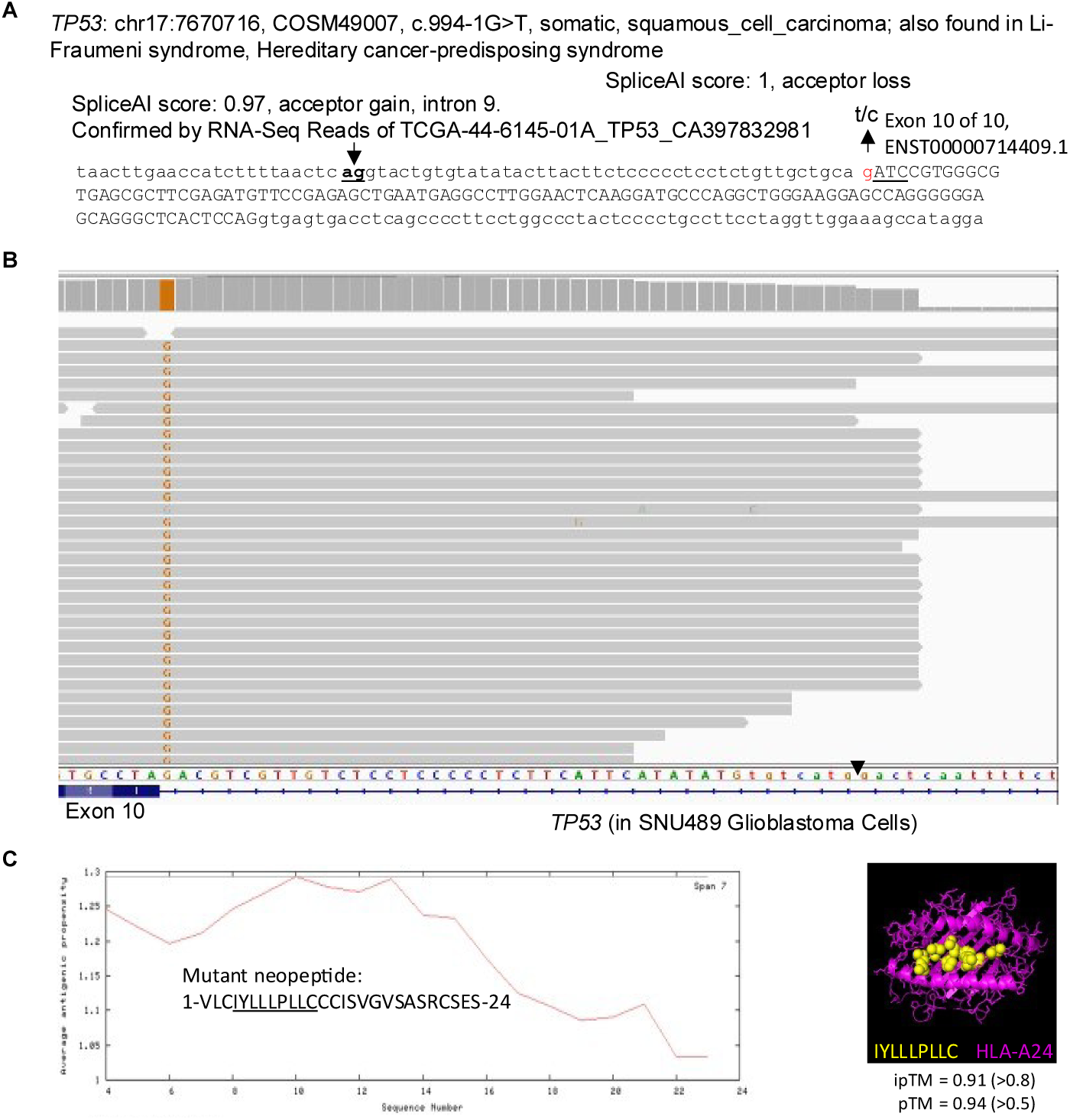
An examples of novel cancer-specific aberrant exon/sequences from SpliceAI prediction of pathogenic SNVs in the intron as a candidate for neo-peptides as cancer-specific neoantigens. **A**. A potential tumor-specific neoantigen from the usage of an intronic cryptic splice site upon the abolishment of the canonical splice site by a SNV. A somatic (or germline in some cases) *TP53* mutation c.994-1, G>T or C, is predicted to cause aberrant splicing and inclusion of an intronic fragment **B**. Evidence of partial intron inclusion by RNA-Seq data from the patient-derived glioblastoma cell line SNU489 carrying the SNV c.994-1, G>C. Arrowhead above the sequence points to the junction between the exon and cryptic splice site AG. **C**. The cancer-specific aberrant splicing-resulted 24aa neopeptide as a potential neoantigen and its predicted interaction with the HLA supertype A24. pTM: confidence for the predicted full structure. ipTM: confidence for the predicted protein-protein interfaces. The bracketed are thresholds for the predicted complex to be similar to the true structure or the confident high-quality predictions, respectively.

To validate the primarily ectopic expression of the aberrant exons in an independent dataset, we filtered against our exome list the reported 10k ‘cancer-specifić exons obtained against a background of 31 normal tissues^29^, which corresponds to 456 tissue pairs in the saturation curve (Fig. 1D). Over 95.8% of them annotated in the hg38 were found in the DEX and 99.9% in our whole exome list. We thus conclude that nearly all aberrant exons are normal tissue exons ectopically expressed in cancer.

### 3. Annotated introns of the normal transcriptome are more likely to contain adenocarcinoma-specific exons

We also examined the annotated introns for aberrantly expressed regions that are potentially specific for cancer, to extend the aberrant target regions further beyond missense mutations and novel exon junctions (neojunctions) in cancer^30-33^.

By pairwise-comparison of the normalized read counts among the 56 tissues by edgeR, we first obtained 329,109 unique differentially expressed intron regions (DEIRs) of 12,975 genes across all chromosomes (S_Fig. 7a-b) as the normal intron background. We then analyzed the intron regions in the adenocarcinomas to obtain 28,007 unique, aberrantly expressed intron regions (AEIR) of 4,448 genes by edgeR-comparison with the corresponding normal tissues of tumor origin (fold change ≥ 4, FDR ≤ 0.01). The AEIR expression increased in 14,523 (52%) regions in at least one of the adenocarcinomas (S_Fig. 8a).

To determine if the increased AEIR in adenocarcinomas are also expressed in any of the normal tissues, we filtered them against the normal transcriptome DEIRs to find that 69% of the increased AEIRs were also among the DEIRs. However, a substantial percentage (31%) of them were highly expressed in the adenocarcinomas (≥ 4 folds) compared to the normal tissues, a much higher abundance than the <1% among the annotated exons. Therefore, the intron regions are more likely to harbor cancer-specific sequences, for which cancer cell-specific single nucleotide variants (SNVs) could be prime candidates for further investigations.

### 4. Authentic cancer-specific splicing events of tumor suppressor genes in association with pathogenic SNVs

To identify authentic cancer-specific exons, particularly those expressed from the intron regions in the 56 tissue background, we examined the aberrant exons and the adenocarcinoma AEIRs for their association with SNVs. Of the SNVs in tumor suppressor genes and oncogenes of 11,070 patients, from the TCGA whole exome sequencing (WES) data in the COSMIC database (Fig. 3A), 3,638 unique SNVs from 6,482 patients were obtained with FATHMM pathogenic score > 0.7, including 1,095 intronic variants with 378 indicated as splice donor/acceptor or splice region variants. We further examined the variants present in our subset of 184 adenocarcinoma patients data obtained through pairing their WES and RNA-Seq data. We identified 57 intron variants from 110 pathogenic ones among 109 patients. Of the variants at the splice sites, cancer-specific AEIRs were observed in the IGV of the corresponding RNA-Seq data for six variants in five genes: *TP53, VHL, PTEN, ACVR1B,* and *SMARCA4*, (S_Table I, S_Fig. 9). Thus, authentic cancer-specific exon sequences are indeed identifiable in the SNV-associated intron or partial intron inclusion events in these patients.

We then used SpliceAI to identify novel splicing-associated genetic variants. For this analysis, we lowered the minimal pathogenic score to 0.5 to cover more variants and analyzed 7085 unique SNVs of the TCGA patients in the COSMIC database. The SpliceAI identified 388 unique variants with confidence scores of higher than 0.5, and 313 unique variants (91% somatic) with highly confident scores of higher than 0.8 (S_Table_II). Their host genes significantly cluster for cancers including breast cancer and leukemia (Fig. 3B, *p* = 5.7E-19 ∼ 5E-9).

In particular, the tumor suppressors *TP53* (95), *PTEN* (18), *RB1* (15), *STK11* (13), *NF1* (9) and *ATM* (8) genes harbor about 50% of the 313 variants, suggesting that tumor suppressor genes are the major targets among the splicing-inducing, pathogenic SNVs in tumorigenesis. Compared to the starting variants of all cancer patients, the SpliceAI not only yielded higher percentages of variants in introns or splice_regions/acceptor/donor sites but also identified a subgroup of variants currently known to cause missense or other non-splicing mutations (Fig. 3C). In total, of the 313 variants with high SpliceAI scores of > 0.8, 53 variants with current mutation description as substitution or deletion/frameshift also created stronger 3’ AG or 5’ GT splice sites; 16 of them had not been known as “splice” variants in genes like *TP53* and *ATM* (S_Table_II). The remaining 260 variants predicted for aberrant splicing are novel with no other known mutation effects yet.

Additionally, we identified 37 variants at the 3’SS together with the (CA)_2_ or G-tract repressor motifs that are prone to cryptic splicing upon disruption by SNVs (S_Table IIIa). The G tracts are enriched in the aberrant 3’SS in cancers^34^. Their 3’SS MaxENT scores are mostly (about 60%) increased in the exons by the SNV alleles, anticipated to promote aberrant splicing. Additionally, we also identified 35 of the pathogenic SNVs to be among the sQTLs of GTEx v10 (S_Table IIIb). Together, the SNV-associated splicing events provide a source for authentic cancer-specific exonic sequences.

An example of predicted effects on cancer-specific aberrant splicing or ATS/APA by representative SNVs is shown in Fig. 3D. Here, deletion of G at the 5’SS of exon 4 of *TP53* (NM_001407264.1) in an invasive breast carcinoma abolished the 5’SS in association with intron retention or APA within intron 4. The latter interpretation is consistent with the declining reads peaks and an AATAA polyA signal motif near the middle of the intron (Fig. 3D). Moreover, reads in the intron’s 3’ half then gradually increased, consistent with an alternative transcription start with the 2^nd^ promoter within the intron^35^, likely resulting in the dominant-negative TP53 isoforms Δ133p53 or Δ160p53 to promote cell growth ^35,36^. In another invasive breast carcinoma, an A>G transition at the upstream 3’SS of the last exon (9) of the *PTEN* gene is associated with the reduced reads of exon 9 and the 3’ half of exon 8, and with alternative transcript termination within intron 8 (S_Fig. 10). The resulting protein would miss amino acids 353-403 at the C-terminal tail, which is more active ^37,38^. More examples are in S_Fig. 11.

Fig. 4 illustrates the potential of such authentic cancer-specific sequences to be translated into novel peptides as potential antigens for cancer vaccine or immunotherapy. The *TP53* somatic mutation, c.994-1,G>T, is predicted by SpliceAI to cause acceptor loss of intron 9 and gain of an upstream cryptic acceptor site (Fig. 4A). Usage of the cryptic site could cause the inclusion of a 44nt intron fragment upstream of the last exon 10 which has been shown by RT-PCR of leukocyte-derived RNA of Li-Fraumeni syndrome family members carrying the T variant^39^. The contribution to the cryptic splice site usage by the T variant is also observed directly in the T-harboring reads of the RNA-Seq data of a TCGA lung adenocarcinoma patient^40^ (S_Fig. 12). Another variant, c.994-1,G>C also causing the acceptor loss, has been found in the human gliobastoma cell line SNU-489^41^. It is the only variant in the reads of the RNA-Seq and the pre-mRNA has nearly 100% cryptic splice site usage (Fig. 4B). The cancer-specific aberrant splicing with partial intron inclusion is expected to result in a 24aa novel peptide in mutation-carriers only. This peptide has high affinity with the HLA-A24 supertype according to the antigen prediction program NetCTL2.0, which are interlocked in the high confidence structural model of the peptide-A24 interaction by AlphaFold3.0 (Fig. 4C).

Taken together, our results support a DEX-dominated human exome. In this complex exome background, nearly all aberrant exons are DEX exons but ectopically expressed in adenocarcinomas. We demonstrate that genuinely cancer-specific exonic sequences can be identified as a result of pathogenic SNV-associated aberrant splicing in introns of tumor suppressor genes, with potential applications as cancer antigens beyond those resulted from point mutations or neojunctions in prevention or immunotherapy.

## Discussion

Humans have the highest percentage of genes (>90%) that have at least one alternative exon, and also have variable exons from alternative transcription start or termination. However, there has not been a clear answer whether it is the differentially expressed exons or the constitutive exons that dominate the transcriptome in the human body. Answers to this question has implications for transcriptome complexity in the presence of the relatively small number of human protein-coding genes, and for the potential development of vaccines against aberrantly expressed antigens such as in cancer. Our data here show that the majority of human exons are DEXs and CEXs are rare. Not surprisingly, nearly all so-called cancer-specific exons in our study are DEXs, so are most of these exons in an independent dataset we examined, emphasizing the need to have sufficient coverage of tissues or development stages to claim the cancer-specificity of an exon. Accordingly, one will need RNASeq data from at least 55 tissues (1485 pairs, > 1463 the threshold) to be able to cover 100% of the DEX exons.

### 1. A DEX-dominating human exome greatly enhances the complexity of the transcriptome

Among the 370,559 substantially detectable exons across 56 types of tissues examined with the current prediction and experimental validation criteria, we estimate that the majority of the exons are DEXs (Fig. 1). This conclusion remains using different thresholds of exon level and fold changes; therefore, the conclusion is likely not a bias due to exon reads levels applied. The high prevalence of DEXs in the human exome is likely still underestimated considering several other factors: (1) The current analysis included only 3 fetal tissues with at least 50 more remaining. (2) Using the rMATS algorithm, which measures reads spanning exon-exon junctions, we identified 50,123 additional alternative splicing events (∼14% more) in 595 tissue pairs that were often not detected by the DEXSeq method, such as the STREX exon ^42,43^. (3) Sex differences were not considered in this analysis, which have been reported in other animal models^44,45^. Thus, it is likely that the majority of human exons are differentially used DEXs among various tissues, developmental stages or sexes and only a smaller proportion of them are truly constitutive exons.

This high percentage of DEXs shifts the paradigm of the human exome from being mainly constitutive to predominantly DEXs, likely to enhance the complexity of the human transcriptome and proteome. Consequently, billions of combinations of variant transcripts or protein isoforms can be generated efficiently and dynamically, contributing to functional diversity and plasticity in human physiology beyond the relatively stable genetic blueprint ^46,47^. Our findings suggest that the definition of many so-called constitutive exons needs to be revisited with more extensive coverage of RNA sources from normal tissues, and so do the mechanisms of their splicing control.

### 2. Cancer specificity of aberrant exons & intron regions depends on the background coverage of normal tissues

Cancer specificity is crucial in diagnosis and therapy. Exploiting the differences between cancer and all noncancerous cells and tissues is thus important particularly in case of precision or personalized medicine treatment. As it is impractical to obtain a wide range of healthy tissues from the same individual, we chose to address this issue by generating a comprehensive catalogue of DEXs or DEIRs of the normal transcriptome as a blueprint background for the adenocarcinomas.

Our cross-checking against the blueprint of 56 normal tissue DEXs revealed that 99% of the aberrant exons of adenocarcinoma are ectopically expressed normal exons, reflecting the highly dynamic spatiotemporal gene regulation. For instance, the E3b-inclusion mediated constitutively active Rac1b variant is likely not only critical for tumorigenesis and metastasis of the lung and colon cancers ^48-52^, but also exists for normal cell physiology in tissues like the esophagus (S_Fig. 5). More importantly, the relative specificity depending on tissue coverage suggests that using corresponding normal tissue as the only control for cancer-specific exons considerably limits our understanding of cancer biology within the human body as a whole. Additionally, our data from the three fetal tissues is consistent with the developmental stage-specific pattern as a splicing dysregulation in cancer.

Several recent studies have reported intron association with disease progression including cancer ^53-55^. Intron retention is one of the mechanisms cancer cells use to inactivate the tumor suppressors and is a potential source of neoepitopes in cancer. Hence, we identified and catalogued the differentially expressed regions of annotated introns across diverse tissues. We also showed that compared to exons, introns possess likely more of the ‘cancer-specifić regions in terms of the number of AEIRs not found among DEIRs of the normal transcriptome. However, more accurate prediction would require full-length transcript sequencing with larger library sizes to minimize artifacts and false-positive results.

Overall, our results highlight the necessity of taking the cancer cells against the whole body transcriptome to identify ‘cancer-specifić exons/intron regions and in addition, to identify the vulnerabilities of the non-cancerous tissues and organs of the body in diagnosis or therapy.

### 3. Integration of associated genetic variants for the identification of cancer-specific exons

Previously it has been reported that the *TP53* transcripts that used the downstream alternative transcription at exon 5, gave rise to N-terminally truncated protein variant Δ133p53 where translation initiated at codon 133 ^56^. The truncated protein variant, lacking the transactivation domain, acts as dominant-negative inhibitors of the wild type p53 ^56^. Our combined analysis of pathogenic variants associated with aberrant intron expression demonstrated that a fragment of intron 4 is expressed in an invasive breast cancer patient from deletion of G at 5’SS of E4 in *TP53* and this is highly specific to the mutant sample, but not in non-mutant or normal breast tissues (Fig. 3B). Together with the genetic variants of *PTEN* (S_Fig. 10) and others (Fig. 3A), these genetic variants provide a rich source for the generation and identification of authentic cancer-specific exons and antigens (Fig. 4) for cancer vaccine or immunotherapy.

## Materials and Methods

### Data Pre-processing

RNA-Seq data of 56 types (53 adult and 3 fetal) of nominally normal human tissues were obtained from the NCBI SRA and dbGaP databases, originally contributed by three sources: 32 adult tissues from a project (PRJEB6971) of the Science for Life Laboratory, Royal Institute of Technology, Stockholm (adipose tissue, adrenal gland, appendix, bone marrow, cerebral cortex, colon, duodenum, endometrium, esophagus, fallopian tube, gallbladder, heart, kidney, liver, lung, lymph node, ovary, pancreas, placenta, prostate, rectum, salivary gland, skeletal muscle, skin, small intestine, smooth muscle, spleen, stomach, testis, thyroid, tonsil, urinary bladder) ^18^; 21 from the Genotype-Tissue Expression (GTEx) (Accession number: phs000424.v7.p2) project of the National Institutes of Health (NIH) (amygdala, anterior cingulate cortex, aorta, breast, caudate, cerebellar hemisphere, cerebellum, cervix, coronary artery, frontal cortex, hippocampus, hypothalamus, nucleus accumbens, pituitary gland, putamen, spinal cord, substantia nigra, tibial artery, tibial nerve, vagina, whole blood) ^19^ and three types of fetal tissues from a project (PRJNA268504) of the Peking University (whole brain, heart and liver) ^20^.

In total, we obtained 12.7 billion of paired-end raw reads (70-100bp) from 433 samples of 56 normal tissues. The quality control measurements were performed by FastQC and Trimmomatic^57^ and we aligned the processed, high-quality reads (35-90bp) to the human transcriptome (Ensembl GRCh38.78) allowing maximum 2 mismatches by TopHat 2.1.1^58^ resulting in 8.8 billion of uniquely mapped reads as concordant pairs for subsequent analyses.

### Annotation Preparation

For the differential exon usage analysis, the reference transcriptome (Ensembl GRCh38.78) was divided into 611,556 bins (hereafter exons) based on the exon boundaries in different transcripts. As the RNA-Seq data used in this study comprised of unstranded reads, hence the exonic regions overlapped between genes were excluded from the reference to minimize the false positive outcomes. Next, we also excluded the exons shorter than 9bp and included the exons with exon base mean (average exon read counts across samples) minimum 20 in any one of the 56 tissue pairs and obtained the reference transcriptome comprising of 370,559 unique exons.

As we know the exon boundaries, using the exon coordinates of Ensembl GRCh38.78, we also divided the introns into bins and obtained a reference transcriptome of the unannotated intragenic regions for the intron differential expression analysis. Introns longer than 200 bp were divided into 100 bp bins.

### Pairwise Analysis of Differential Exon Usage

We have used DEXSeq ^23^ for differential exon usage analysis across tissues where the ratio of read counts of the target exon to the read counts of all other exons of the same gene infers differentially used exons (DEXs) including both internal and terminal exons. Therefore, DEXSeq identifies DEXs mediated by alternative splicing (AS), alternative transcription start (ATS) or alternative polyadenylation (APA). In order to identify the maximum number of DEXs across tissues, we have compared the tissues pairwise in all possible combinations (C(56,2)) comprising 1,540 tissue-pairs in total.

We have used stringent filtering criteria as reported in our previous study ^24^ to obtain a confident list of DEXs where exons with exon base mean (EBM) less than 20 (the mean of read counts across all tissue pairs for that exon), average base mean (ABM) less than 20 (the mean of read counts across all tissue pairs for that gene), the ratio of target exon-EBM (normalized to the exon length) to the highest EBM in that gene, less than 0.002, adjusted *p* value higher than 0.05 and fold change less than 0.1 were excluded.

In addition to DEXSeq, for a group of tissues (35) we have also used replicate MATS (Multivariate Analysis of Transcript Splicing) ^59^ that provided additional splicing events (FDR < 0.05) not overlapped with the DEXSeq outputs.

### Tagging Terminal Exons

To estimate the relative contribution by each of AS, ATS and APA mediating DEXs, we have tagged all DEXs as whether or not terminal exons using the coordinates of different transcripts for each host gene. Since the DEXs co-mediated by AS and ATS/APA could be very complex to comprehend, we grouped the terminal DEXs considering the comparatively simpler situations as follows -

DEXs containing different transcription start sites with the same splice donor sites

DEXs containing different polyadenylation sites with the same splice acceptor sites

DEXs containing alternative splice sites with the same transcription start or polyadenylation sites

### Pairwise Analysis of Intron Differential Expression

Using our transcriptome reference of the unannotated regions, the reads mapped to intron bins were counted by FeatureCounts^60^. The intron read counts normalized to their gene level were compared between tissues by edgeR ^61^ to identify the differentially expressed intron regions (DEIRs) across tissues. We have excluded the intron regions shared by multiple genes of the same or opposite strand and further shortlisted the differentially expressed intron regions (DEIRs) with FDR ≤ 0.01 and a minimum of Log2 fold changes. Similarly as exons, the intron analyses were also performed pairwise in 1540 tissue-pairs from 56 tissue-types.

### Validation by RT-PCR after DEXSeq Analysis

We validated 244 out of 269 alternative splicing events from 24 exons in total by semi-quantitative RT-PCR: 10 in sixteen human tissues (adrenal gland, bone marrow, brain, colon, fetal brain, fetal liver, liver, kidney, lung, placenta, prostate, salivary gland, skeletal muscle, spleen, small intestine and testis), and 14 in GH3 cells from a separate study of the same DEXSeq filtering criteria. Human tissue RNA samples (1µg/µl) were purchased from Clontech Laboratories Inc. (Cat No. 636643)^62^. About 500ng of RNA was used for 20 µl of reverse transcription reaction. Polymerase Chain Reaction (PCR) for splicing analysis was performed with 30 cycles for different genes. PCR products were resolved in 2.5-3.5% agarose gels containing ethidium bromide and visualized with a digital camera under ultraviolet light. The percentages of exon inclusion were calculated from the band intensities and normalized by their product lengths to molar ratio.

### Functional Annotation Clustering

Functional annotation and classification of related genes were analysed using DAVID under high stringent conditions ^63^.

### Analysis of Adenocarcinoma Data

The RNA-Seq data of 184 patients from five types of adenocarcinomas: lung (54 patients), colon (15), pancreatic (35), prostate (36) and invasive breast carcinoma (44) were downloaded from the Cancer Genome Atlas (TCGA) database (Accession number: phs000178.v9.p8) ^27^ ^28^. The raw data (50bp long paired-end reads) were processed similarly as the normal tissues described above. The samples of adenocarcinoma for each type were compared with their corresponding normal by DEXSeq and edgeR to identify the aberrantly expressed exons and introns in adenocarcinoma, respectively.

### Validation of Aberrant Splicing by Semi-quantitative RT-PCR

The tumor samples were purchased from the Ontario Tumor Bank. For RNA extraction, the frozen tumor samples were excised to get tissue from the tumor centre and homogenized immediately with appropriate volume of lysis buffer (with β-ME) and step-by-step tissue RNA purification protocol for Qiagen RNeasy mini kits was followed. The predicted aberrant splicing events from the above TCGA RNA-Seq analysis were validated using the extracted RNAs from the tumor samples by semi-quantitative RT-PCR similarly as described above for the normal tissues.

### Determining the antigenicity of peptide sequence

The nucleotide sequences of the identified regions were translated into protein/peptide sequences in frame with the upstream coding reigons. The antigenicity of the peptide regions were determined at the webserver http://imed.med.ucm.es/Tools/antigenic.pl ^64^. The CTL epitopes of the protein/peptide sequences were determined using NetCTL 2.0 ^65^. IGV verification also confirmed that epitopes were associated with regions highly expressed in the RNA-seq samples of cancer patients or patient-derived cell lines carrying the genetic variants. The antigenic neopeptide interaction with the HLA supertype A24 was modeled in Google AlphaFold3.0 with high confidence scores >0.9 ^66^.

## Acknowledgements

We would like to thank the NCBI dbGaP staffs, GTEx and <u>TCGA research network</u>, the NIH data repository, and the contributing investigators for making the data available. Special thanks to the U of M Med IT team, particularly Nelson Vieira and Gilles Detillieux for their help with the data management process of this study. The study has been supported in part by Discovery Grants from the Natural Sciences and Engineering Research Council of Canada (NSERC, RGPIN-2016-06004 and RGPIN-2022-05023), and a Manitoba Research Chair fund to J.X.; by a UMGF scholarship to U.D. and Research Manitoba Graduate Studentship to H.N.

## References

1 Kahles, A. et al. Comprehensive Analysis of Alternative Splicing Across Tumors from 8,705 Patients. Cancer Cell 34, 211–224 e216, doi:10.1016/j.ccell.2018.07.001 (2018).

2 David, J. K., Maden, S. K., Weeder, B. R., Thompson, R. F. & Nellore, A. Putatively cancer-specific exon-exon junctions are shared across patients and present in developmental and other non-cancer cells. NAR Cancer 2, zcaa001, doi:10.1093/narcan/zcaa001 (2020).

3 Oltean, S. & Bates, D. O. Hallmarks of alternative splicing in cancer. Oncogene 33, 5311–5318, doi:10.1038/onc.2013.533 (2014).

4 Wiesner, T. et al. Alternative transcription initiation leads to expression of a novel ALK isoform in cancer. Nature 526, 453–457, doi:10.1038/nature15258 (2015).

5 Roudnicky, F. et al. Alternative transcription of a shorter, non-anti-angiogenic thrombospondin-2 variant in cancer-associated blood vessels. Oncogene 37, 2573–2585, doi:10.1038/s41388-018-0129-z (2018).

6 Xiang, Y. et al. Comprehensive Characterization of Alternative Polyadenylation in Human Cancer. J Natl Cancer Inst 110, 379–389, doi:10.1093/jnci/djx223 (2018).

7 Nilsen, T. W. & Graveley, B. R. Expansion of the eukaryotic proteome by alternative splicing. Nature 463, 457–463, doi:10.1038/nature08909 (2010).

8 Kim, E., Magen, A. & Ast, G. Different levels of alternative splicing among eukaryotes. Nucleic acids research 35, 125–131, doi:10.1093/nar/gkl924 (2007).

9 Maniatis, T. & Tasic, B. Alternative pre-mRNA splicing and proteome expansion in metazoans. Nature 418, 236–243, doi:10.1038/418236a (2002).

10 Reyes, A. & Huber, W. Alternative start and termination sites of transcription drive most transcript isoform differences across human tissues. Nucleic acids research 46, 582–592, doi:10.1093/nar/gkx1165 (2018).

11 Shabalina, S. A., Ogurtsov, A. Y., Spiridonov, N. A. & Koonin, E. V. Vol 42 7132-7144 (2014).

12 Pan, Q., Shai, O., Lee, L. J., Frey, B. J. & Blencowe, B. J. Deep surveying of alternative splicing complexity in the human transcriptome by high-throughput sequencing. Nat Genet 40, 1413–1415 (2008).

13 Wang, E. T. et al. Alternative isoform regulation in human tissue transcriptomes. Nature 456, 470–476, doi:10.1038/nature07509 nature07509 [pii] (2008).

14 Jayasinghe, R. G. et al. Systematic Analysis of Splice-Site-Creating Mutations in Cancer. Cell Rep 23, 270–281 e273, doi:10.1016/j.celrep.2018.03.052 (2018).

15 Jung, H., Lee, K. S. & Choi, J. K. Comprehensive characterisation of intronic mis-splicing mutations in human cancers. Oncogene 40, 1347–1361, doi:10.1038/s41388-020-01614-3 (2021).

16 Cao, S. et al. Discovery of driver non-coding splice-site-creating mutations in cancer. Nat Commun 11, 5573, doi:10.1038/s41467-020-19307-6 (2020).

17 Dentro, S. C. et al. Characterizing genetic intra-tumor heterogeneity across 2,658 human cancer genomes. Cell 184, 2239-+, doi:10.1016/j.cell.2021.03.009 (2021).

18 Uhlen, M. et al. Proteomics. Tissue-based map of the human proteome. Science 347, 1260419, doi:10.1126/science.1260419 (2015).

19 Consortium, G. T. The Genotype-Tissue Expression (GTEx) project. Nat Genet 45, 580–585, doi:10.1038/ng.2653 (2013).

20 Yan, L. et al. Epigenomic Landscape of Human Fetal Brain, Heart, and Liver. J Biol Chem 291, 4386–4398, doi:10.1074/jbc.M115.672931 (2016).

21 Zhang, J. & Manley, J. L. Misregulation of pre-mRNA alternative splicing in cancer. Cancer discovery 3, 1228–1237, doi:10.1158/2159-8290.CD-13-0253 (2013).

22 Feng, D. & Xie, J. Aberrant splicing in neurological diseases. Wiley interdisciplinary reviews. RNA 4, 631–649, doi:10.1002/wrna.1184 (2013).

23 Anders, S., Reyes, A. & Huber, W. Detecting differential usage of exons from RNA-seq data. Genome research 22, 2008–2017, doi:10.1101/gr.133744.111 (2012).

24 Lei, L. et al. Multilevel Differential Control of Hormone Gene Expression Programs by hnRNP L and LL in Pituitary Cells. Mol Cell Biol 38, doi:10.1128/MCB.00651-17 (2018).

25 Robinson, J. T. et al. Integrative genomics viewer. Nat Biotechnol 29, 24–26, doi:10.1038/nbt.1754 (2011).

26 Sung, H. et al. Global cancer statistics 2020: GLOBOCAN estimates of incidence and mortality worldwide for 36 cancers in 185 countries. Ca-Cancer J Clin 71, 209–249, doi:10.3322/caac.21660 (2021).

27 Weinstein, J. N. et al. The Cancer Genome Atlas Pan-Cancer analysis project. Nature Genetics 45, 1113–1120, doi:10.1038/ng.2764 (2013).

28 Schlomm, T. Results of the CGC/TCGA Pan-Cancer Analysis of the Whole Genomes (PCAWG) Consortium. Urologe 59, 1552–1553, doi:10.1007/s00120-020-01373-9 (2020).

29 Wu, S. et al. ASCancer Atlas: a comprehensive knowledgebase of alternative splicing in human cancers. Nucleic acids research 51, D1196–D1204, doi:10.1093/nar/gkac955 (2023).

30 Kwok, S. & Higuchi, R. Avoiding false positives with PCR. Nature 339, 237–238 (1989).

31 Wu, D., Gallagher, D. T., Gowthaman, R., Pierce, B. G. & Mariuzza, R. A. Structural basis for oligoclonal T cell recognition of a shared p53 cancer neoantigen. Nat Commun 11, 2908, doi:10.1038/s41467-020-16755-y (2020).

32 Hsiue, E. H. et al. Targeting a neoantigen derived from a common TP53 mutation. Science 371, doi:10.1126/science.abc8697 (2021).

33 Malekzadeh, P. et al. Neoantigen screening identifies broad TP53 mutant immunogenicity in patients with epithelial cancers. J Clin Invest 129, 1109–1114, doi:10.1172/JCI123791 (2019).

34 Nguyen, H. & Xie, J. Widespread Separation of the Polypyrimidine Tract From 3’ AG by G Tracts in Association With Alternative Exons in Metazoa and Plants. Front Genet 9, 741, doi:10.3389/fgene.2018.00741 (2018).

35 Bourdon, J. C. et al. p53 isoforms can regulate p53 transcriptional activity. Genes Dev 19, 2122–2137, doi:10.1101/gad.1339905 (2005).

36 Marcel, V. et al. Delta160p53 is a novel N-terminal p53 isoform encoded by Delta133p53 transcript. FEBS Lett 584, 4463–4468, doi:10.1016/j.febslet.2010.10.005 (2010).

37 Mingo, J. et al. Precise definition of PTEN C-terminal epitopes and its implications in clinical oncology. NPJ Precis Oncol 3, 11, doi:10.1038/s41698-019-0083-4 (2019).

38 Vazquez, F., Ramaswamy, S., Nakamura, N. & Sellers, W. R. Phosphorylation of the PTEN tail regulates protein stability and function. Mol Cell Biol 20, 5010–5018, doi:10.1128/MCB.20.14.5010-5018.2000 (2000).

39 Verselis, S. J., Rheinwald, J. G., Fraumeni, J. F., Jr. & Li, F. P. Novel p53 splice site mutations in three families with Li-Fraumeni syndrome. Oncogene 19, 4230–4235, doi:10.1038/sj.onc.1203758 (2000).

40 Palmisano, A., Vural, S., Zhao, Y. & Sonkin, D. MutSpliceDB: A database of splice sites variants with RNA-seq based evidence on effects on splicing. Hum Mutat 42, 342–345, doi:10.1002/humu.24185 (2021).

41 Shin, K. H. et al. Establishment and characterization of nine human brain tumor cell lines. In Vitro Cell Dev Biol Anim 37, 625–628, doi:10.1290/1071-2690(2001)037<0625:EACONH>2.0.CO;2 (2001).

42 Xie, J. Y. & McCobb, D. P. Control of alternative splicing of potassium channels by stress hormones. Science 280, 443–446, doi:DOI 10.1126/science.280.5362.443 (1998).

43 Xie, J. & Black, D. L. A CaMK IV responsive RNA element mediates depolarization-induced alternative splicing of ion channels. Nature 410, 936 (2001).

44 Ray, M. et al. Sex-specific splicing occurs genome-wide during early Drosophila embryogenesis. Elife 12, doi:10.7554/eLife.87865 (2023).

45 Rogers, T. F., Palmer, D. H. & Wright, A. E. Sex-Specific Selection Drives the Evolution of Alternative Splicing in Birds. Mol Biol Evol 38, 519–530, doi:10.1093/molbev/msaa242 (2021).

46 Razanau, A. & Xie, J. Y. Emerging mechanisms and consequences of calcium regulation of alternative splicing in neurons and endocrine cells. Cell Mol Life Sci 70, 4527–4536, doi:10.1007/s00018-013-1390-5 (2013).

47 Baralle, F. E. & Giudice, J. Alternative splicing as a regulator of development and tissue identity. Nat Rev Mol Cell Bio 18, 437–451, doi:10.1038/nrm.2017.27 (2017).

48 Jordan, P., Brazao, R., Boavida, M. G., Gespach, C. & Chastre, E. Cloning of a novel human Rac1b splice variant with increased expression in colorectal tumors. Oncogene 18, 6835–6839, doi:10.1038/sj.onc.1203233 (1999).

49 Fiegen, D. et al. Alternative splicing of Rac1 generates Rac1b, a self-activating GTPase. J Biol Chem 279, 4743–4749, doi:10.1074/jbc.M310281200 (2004).

50 Radisky, D. C. et al. Rac1b and reactive oxygen species mediate MMP-3-induced EMT and genomic instability. Nature 436, 123–127, doi:10.1038/nature03688 (2005).

51 Kotelevets, L. & Chastre, E. Rac1 Signaling: From Intestinal Homeostasis to Colorectal Cancer Metastasis. Cancers (Basel) 12, doi:10.3390/cancers12030665 (2020).

52 Fu, X. D. Both sides of the same coin: Rac1 splicing regulating by EGF signaling. Cell Res 27, 455–456, doi:10.1038/cr.2017.19 (2017).

53 Flodrops, M. et al. TIMP1 intron 3 retention is a marker of colon cancer progression controlled by hnRNPA1. Mol Biol Rep 47, 3031–3040, doi:10.1007/s11033-020-05375-w (2020).

54 Zhang, D. X. et al. Intron retention is a hallmark and spliceosome represents a therapeutic vulnerability in aggressive prostate cancer. Nature Communications 11, doi:ARTN 2089 10.1038/s41467-020-15815-7 (2020).

55 Tan, D. J., Mitra, M., Chiu, A. M. & Coller, H. A. Intron retention is a robust marker of intertumoral heterogeneity in pancreatic ductal adenocarcinoma. Npj Genom Med 5, doi:ARTN 55 10.1038/s41525-020-00159-4 (2020).

56 Courtois, S. et al. DeltaN-p53, a natural isoform of p53 lacking the first transactivation domain, counteracts growth suppression by wild-type p53. Oncogene 21, 6722–6728, doi:10.1038/sj.onc.1205874 (2002).

57 Bolger, A. M., Lohse, M. & Usadel, B. Trimmomatic: a flexible trimmer for Illumina sequence data. Bioinformatics 30, 2114–2120, doi:10.1093/bioinformatics/btu170 (2014).

58 Kim, D. et al. TopHat2: accurate alignment of transcriptomes in the presence of insertions, deletions and gene fusions. Genome Biol 14, R36, doi:10.1186/gb-2013-14-4-r36 (2013).

59 Shen, S. et al. rMATS: robust and flexible detection of differential alternative splicing from replicate RNA-Seq data. Proc Natl Acad Sci U S A 111, E5593–5601, doi:10.1073/pnas.1419161111 (2014).

60 Liao, Y., Smyth, G. K. & Shi, W. featureCounts: an efficient general purpose program for assigning sequence reads to genomic features. Bioinformatics 30, 923–930, doi:10.1093/bioinformatics/btt656 (2014).

61 Robinson, M. D., McCarthy, D. J. & Smyth, G. K. edgeR: a Bioconductor package for differential expression analysis of digital gene expression data. Bioinformatics 26, 139–140, doi:10.1093/bioinformatics/btp616 (2010).

62 Sohail, M. et al. Differential expression, distinct localization and opposite effect on Golgi structure and cell differentiation by a novel splice variant of human PRMT5. Biochim Biophys Acta 1853, 2444–2452, doi:10.1016/j.bbamcr.2015.07.003 (2015).

63 Huang da, W., Sherman, B. T. & Lempicki, R. A. Systematic and integrative analysis of large gene lists using DAVID bioinformatics resources. Nat Protoc 4, 44–57, doi:10.1038/nprot.2008.211 (2009).

64 Kolaskar, A. S. & Tongaonkar, P. C. A semi-empirical method for prediction of antigenic determinants on protein antigens. FEBS Lett 276, 172–174, doi:10.1016/0014-5793(90)80535-q (1990).

65 Larsen, M. V. et al. Large-scale validation of methods for cytotoxic T-lymphocyte epitope prediction. BMC Bioinformatics 8, 424, doi:10.1186/1471-2105-8-424 (2007).

66 Abramson, J. et al. Accurate structure prediction of biomolecular interactions with AlphaFold 3. Nature 630, 493–500, doi:10.1038/s41586-024-07487-w (2024).

